# Fructose malabsorption induces dysbiosis and increases anxiety in Human and animal models

**DOI:** 10.1101/2025.03.25.645339

**Authors:** Adeline Coursan, Delphine Polve, Anne-Marie Leroi, Magali Monnoye, Lea Roussin, Marie-Pierre Tavolacci, Muriel Quillard Muraine, Mathilde Maccarone, Olivia Guérin, Estelle Houivet, Charlène Guérin, Valery Brunel, Jérôme Bellenger, Jean-Paul Pais de Barros, Guillaume Gourcerol, Laurent Naudon, Sophie Layé, Charlotte Madore, Xavier Fioramonti, Chloé Melchior, Véronique Douard

## Abstract

**Background & Aims:** Excessive fructose intake is a growing public health concern, yet many individuals have a lower absorption capacity than the average intake, leading to widespread chronic fructose malabsorption. This results in intestinal fructose spillover, disrupting gut microbiota and triggering peripheral inflammation, which, along with neuroinflammation, plays a key role in mood disorders. This study investigates the connection between fructose malabsorption and mood disorders by examining gut microbiota changes in a human cohort and exploring their links with neuroinflammation in a GLUT5-KO mouse model.

**Methods:** In a human cohort, fructose malabsorption was assessed using a breath hydrogen test, while plasma lipopolysaccharide (LPS) levels and anxiety traits (measured using the State-Trait Anxiety Inventory, STAI) were analyzed. Gut microbiota composition was characterized through 16S rRNA sequencing, and dietary fructose intake was recorded. In the preclinical study, Glut5-KO mice, which lack intestinal fructose transport, were fed a 5% fructose diet for four weeks. Behavioral assays assessed anxiety- and depressive-like behaviors, while gut microbiota composition and microglia-associated gene expression were analyzed.

**Results:** Among the recruited healthy volunteers, 60% exhibited fructose malabsorption, along with elevated plasma LPS levels, increased anxiety traits on the STAI, and distinct gut microbiota alterations, partially linked to fructose intake patterns. The average daily fructose intake was 30 g per individual, with significant variability in dietary sources. In the preclinical model, Glut5-KO mice on a 5% fructose diet displayed increased anxiety- and depressive-like behaviors, pronounced gut microbiota shifts, and altered expression of microglia-associated genes.

**Conclusions:** These findings highlight the complex interplay between dietary fructose, gut microbiota, and neuroinflammation in shaping mental health. Chronic fructose malabsorption may contribute to mood disorders through gut dysbiosis and microglia-dependent neuroinflammation, warranting further investigation into dietary interventions.

**HIGHLIGHTS:**

- Fructose malabsorption is associated with anxiety traits in healthy volunteers.
- Fructose malabsorption enhances anxiety-like behaviors in malabsorptive Glut5-KO mice.
- Fructose malabsorption is associated with gut microbiota dysbiosis in human and preclinical mouse model of fructose malabsorption in association with fructose intake
- Fructose malabsorption increases neuroinflammation and alters microglia functions in malabsorptive Glut5-KO mice.

## BACKGROUND

The maintenance of brain functions is intricately linked to dietary factors and the impact of poor nutrition on cerebral functions has emerged as a significant public health concern [1, 2]. Several studies have highlighted the role of dietary carbohydrates in neurological functions and related disorders including mood disorders [3–6]. Among these carbohydrates, fructose is of particular interest, as various international public health organizations have highlighted its overconsumption as a potential threat to human health [7, 8]. Historically, humans consumed less than 5 grams of fructose daily for thousands of years, whereas current consumption levels range from 50 to 80 grams per day in developed countries [9]. Fructose is primarily absorbed via the GLUT5 transporter that is up-regulated by luminal fructose in a ketohexokinase (KHK)- dependent manner [10]. However, GLUT5 is saturable leading to a limited capacity of the small intestine to fully absorb fructose, causing a widespread chronic malabsorption in the adult population in which up to 60% of healthy adult could not absorbed a load of 40 grams of fructose [11, 12]. Consequently, fructose intake can result in its spillover into the distal parts of the intestine, where it interacts with colonic bacteria, generating degradation products such as short-chain fatty acids (SCFA), carbon dioxide (CO₂), hydrogen (H₂), or methane (CH₄) [12, 13]. In addition, incomplete absorption of fructose also alter gut physiology and major changes in microbiota composition, as demonstrated in animal models of fructose malabsorption [14, 15].

Both human and animal studies have shown that changes in microbiota composition can significantly impact brain health, including increasing the risk of anxiety or depressive symptoms [16–18]. Changes in microbiota composition and metabolism have been associated with increased neuroinflammation through different mechanisms involving the release of microbiota-derived molecules or enhancement of peripheral inflammation, which can both, impact the brain and promote neuroinflammation [16, 19]. Neuroinflammation plays a crucial role in the development of mood disorders, including anxiety and depressive-like behaviors [17, 20, 21]. It is a protective immune response ensured by microglia—the resident immune cells of the brain [22]— which are especially important in these processes, with gut microbiota metabolites such as SCFAs or microbiota-derived indoles playing a key role in regulating microglia homeostasis in mice [23–27]. High-fructose feeding has been associated with neuroinflammation and brain dysfunctions including anxiety- and depressive-like behaviors in rodent models [28–31]. Interestingly, a higher prevalence of anxiety has also been observed in individuals with carbohydrate malabsorption [32, 33], including fructose malabsorption [34]. However, the mechanisms underlying this association remain unknown. Therefore, we hypothesized that fructose malabsorption increases the risk of mood disorders, such as anxiety, by altering gut microbiota composition, which will subsequently impair microglia homeostasis and promote neuroinflammation. To test the hypothesis that fructose malabsorption is linked to anxiety, we first compared anxiety traits in healthy volunteers with and without fructose malabsorption using the MoodyFructose cohort, which was specifically developed for this study. We also determined their fructose consumption profiles, measure specific biochemical markers of inflammation and analyzed their gut microbiota composition. Additionally, we developed correlations between microbiota composition, biochemical markers, anxiety scores, and dietary patterns to identify potential mechanisms underlying these relationships. Finally, we examined emotional behaviors, gut microbiota composition, and microglial gene expression in GLUT5-KO mice, a preclinical model for fructose malabsorption, to further investigate the potential mechanisms linking fructose malabsorption to mood disorders.

## MATERIALS AND METHODS

### Human study

#### Study population

Moodyfructose was approved by the national ethical committee “Comité de protection des personnes Ouest II*”* on 25^th^ of June 2021 and was registered at www.clinicaltrials.gov (NCT05371067). Fifty-five volunteers gave written informed consent before participating. The study protocol conformed to the ethical guidelines of the 1975 Declaration of Helsinki. The study (MoodyFructose)’s primary aim was to compare anxiety traits in healthy volunteers with and without fructose malabsorption. To specifically target a population with high fructose intake [35, 36], we included only males aged 18 to 35 with a body mass index (BMI) below 25. This criterion helped minimize confounding effects associated with overweight and obesity [37] ensuring a homogeneous study group with consistent metabolic profiles. Exclusion criteria included the presence of gastrointestinal disorders (including irritable bowel syndrome, celiac disease, and inflammatory bowel disorders), psychiatric conditions, neurological diseases, eating disorders, diabetes, a history of digestive surgery, or any chronic disease or current infectious disease (Supp Table S1). The volunteers were not allowed to smoke, undergo long-term treatment, follow any specific diet, or have taken antibiotics in the last six months or probiotics in the last three months. Volunteers were excluded if they had a positive glucose breath test, did not follow the recommendations before the breath tests (Supp Table S1), and if the State-Trait Anxiety Inventory (STAI) questionnaire was not performed at baseline.

#### Breath tests

Each volunteer underwent two breath tests, one during the second visit and one during the last visit, within 8 to 31 days after inclusion. These tests were conducted according to the European guidelines using the same preparation and methods as previously published [38]. Briefly, Patients followed a strict preparation, including a specific diet for 48 hours before the breath tests [39]. Then, the fructose breath test was performed with a 35g fructose load [38]. End-alveolar breath samples were collected every 30 min for five hours during the fructose breath test. Both H2 and CH4 levels were measured and the test was considered positive for fructose malabsorption when H2 and/or CH4 levels rose above 20 ppm. To ensure the specificity of the test and rule out small intestinal bacterial overgrowth, which can lead to false-positive results in the fructose breath test, a glucose breath test was performed using a 75g glucose load [39, 40]. Following European guidelines, end-alveolar breath samples of expired air were collected every 15 min for two hours and the test was considered positive if any of the following occurred: a rise of H2 or CH4 levels above 20 ppm from baseline, or an increase in H2 or CH4 levels above 10 ppm in two consecutive samples compared to individual baseline levels [39]. Volunteers with a positive glucose breath test were excluded from the study.

#### Fructose intake assessment

Volunteers were asked to complete a 7-day food diary at home, documenting all food sources consumed between the first and final visit to assess their fructose intake profile. The dietary surveys were based on a meal-centered approach, with questions about the types and quantities of food consumed during meals or snacking. For each food item, the total quantity of carbohydrate, fructose and sucrose was obtained using the Ciqual food composition database developed by the French Agency for Food, Environmental and Occupational Health & Safety (ANSES) (https://ciqual.anses.fr/). Added sugars were defined as all monosaccharides and disaccharides added to foods and beverages by the manufacturer, cook or consumer, and sugars naturally present in honey, syrups, fruit juices and fruit juice concentrates. Sweet products included cakes, pastries, biscuits, Viennese pastries, confectioneries, honey, marmalade, chocolate spread and sweet dairies. Beverages included fruit juices, sodas, smoothies, chocolate milk, alcohol and sweet coffee or tea. Fruits and vegetables include fresh and processed fruits and vegetables as well as dry fruits or nuts. Dairy included cheese, plain yogurt and plain milk. Cereals include oats, wheat, barley, rye, quinoa and whole-grain bread.

#### Anxiety and depression questionnaires

The STAI and the Hospital Anxiety and Depression Scale (HADS) questionnaires were completed by all volunteers at baseline and during the second visit. STAI assesses anxiety traits, including feelings of apprehension, tension, nervousness, and concern [41]. It is considered an indicator of transient changes in anxiety due to aversive situations. The questionnaire consists of 20 items, each rated on a 4-point Likert scale (1 = not at all, 4 = very much so), with total scores ranging from 20 to 80. Higher scores indicate greater anxiety symptoms, with a score above 37 suggesting a state of anxiety [42]. The HADS evaluates pathological anxiety and depression and was used in this study potentially identify individuals with severe symptoms. It consists of 14 items, divided into two subscales: anxiety (7 items) and depression (7 items). Each item is scored from 0 to 3 [43], with higher scores indicating more severe psychological distress. Each subscale ranges from 0 to 21, with scores above 11 indicating clinical anxiety or depression.

#### Human Samples and analysis

Stool samples were collected during the final visit, sampled and stored at the Centre de Ressources Biologiques (CIC-CRB U1404, Rouen, France). Briefly, the stools were aliquoted into microtubes without chemical or biological additives and frozen at −80°C within 6 hours of release and remained frozen until microbiota analysis following gold standard procedures [44]. For calprotectin analysis, non-frozen stools (before storage at −80C) were used and extracted using CALEX®-cap devices and calprotectin levels were determined by an enzyme-linked immunosorbent assay (ELISA) (Bühlmann Labs, Schönenbuch, Switzerland) on 50– 150 µL of extracts as described previously [45]. Blood was sampled from fasting volunteers during the final visit (before breath tests) to analyze C-reactive protein (CRP) levels and lipopolysaccharides (LPS). CRP levels were measured using a Cobas® 8000 chemistry analyzer (Roche Diagnostics, Mannheim, Germany) in the biochemistry laboratory of the CIC-CRB (Rouen, France). Plasma LPS concentrations were determined by high-performance liquid chromatography coupled with gas-phase/tandem mass spectrometry, as previously described [46].

##### Animals

Adult WT (n = 8 to 10) or Glut5_KO (n = 8 to 10) male mice (8-12 weeks) were fed with a 5% fructose diet (% from kcal) (Research diet, New Jersey, USA) for 4 weeks and group-housed in conventional housing cages at Infectiologie Expérimentale des Rongeurs et Poissons (IERP, Jouy-en-Josas) animal facility and at the NutriNeuro animal facility (CIRCE Bordeaux). A group of Glut5_KO male mice of the same age were fed with a 0 % fructose diet (0 % fructose diet was isocaloric and content the same amount of carbohydrate than the 5 % fructose diet; Table 1). Animals were maintained on a 12-hours light-dark cycle with *ad libitum* access to food and water. All experimental procedures were conducted in accordance with the European directive 2010/63/UE and approved by the French Ministry of Research and local ethics committees (APAFIS#: 19870 and APAFIS#23075).

**Table 1:** Diets composition.

| DIET COMPOSITION | 0% FRUCTOSE | 5% FRUCTOSE |
| --- | --- | --- |
| **PROTEINS (KCAL%)** | **20** | **20** |
| **LIPIDS (KCAL%)** | **16** | **16** |
| **CARBOHYDRATES (KCAL%)** | **64** | **64** |
| Corn starch (gm%) | 329 | 329 |
| Sucrose (gm%) | 0 | 0 |
| Dextrose (gm%) | 200 | 150 |
| Fructose (gm%) | 0 | 50 |
| Cellulose BW2000 (gm%) | 50 | 50 |
| **Total kcal/kg** | **3964** | **3964** |

##### Behavioral tests

Behavioral tests were performed after 4-weeks of diet while animals (n = 8 per group) were maintained under the same diet during the tests. Animals were tested in different anxiety-like and depressive-like behavioral paradigms as previously described [47–50]. Briefly, the elevated plus maze (EPM) test was performed for 5 minutes in an apparatus elevated to 120 cm from the floor, including two 35 cm length and 5cm wide open arms and two closed arms with same dimensions and surrounded with 15 cm high wall. The forced swim test (FST) was carried out for 6 minutes using a glass cylinder tank filled with water (26 ± 1 °C) in a manner that the mouse cannot reach the floor with its tail. Subsequently, the total immobility time of the mice in the water during the 6 minutes of the test was recorded. The tests were spaced by two days of recovery time and increasing in term of stress-related paradigm, starting with the less stressful test (EPM). All videos were analyzed using the ANY-maze software (Stoelting Co., Dublin, Ireland), automatically count on the software in a blinded setting.

##### Microglia gene expression

Microglia isolation was performed as previously described [51]. Briefly, mice (were euthanized by intraperitoneal pentobarbital injection (Exagon®, 300 mg/kg), 30 min post-buprenorphine administration (0.1 mg/kg) before being transcardially perfused with ice-cold Hanks’ Balanced Salt Solution (HBSS, Gibco). Following brain’s mechanical dissociation, single cell suspensions were prepared and centrifuged over a 37%/70% discontinuous Percoll gradient (GE Healthcare), mononuclear cells were isolated from the interphase. Total RNAs were extracted using RNeasy Plus Mini Kit (Qiagen, 74106) according to the manufacturer’s protocol. RNA concentration and purity were determined using Nanodrop spectrophotometer (Nanodrop technologies). 10 µL of RNAs were reverse transcribed using Superscript IV VILO (Invitrogen, Life Technologies, France). cDNAs were amplified using commercially available FAM-labeled Taqman® probes (Applied Biosystems), [*glut5* (Mm00600311_m1), *csf1r* (Mm01266652_m1), *p2ry12* (Mm00446026_m1), *clec7a* (Mm01183349_m1), *spp1* (Mm00436767_m1), *apoe* (Mm01307193_g1), *il-1b* (Mm00434228_m1), *il-6* (Mm00446190_m1), *tnf-a* (Mm00443258_m1), *tgfbr2* (Mm03024091_m1), *B2m* (Mm00437762_m1, used as reference gene)]. Real-time PCR reaction was performed in duplicate using a LightCycler® 480 instrument II (Roche). Gene expression data are normalized relative to B2m expression and to the control group GLUT5_KO5F (2^(−ΔΔCt)).

##### Microbiota composition

Total DNA was extracted from mice cecal content and healthy volunteers’ feces samples with PowerFecal Pro DNA Kit (Qiagen, France) and using the mechanical bead-beating disruption method. The V3–V4 hypervariable region of the bacterial 16S rDNA was amplified by PCR (PCR1F_460: CTTTCCCTACACGACGCTCTTCCGATCTACGGRAGGCAGCAG and PCR1R_460: GGAGTTCAGACGTGTGCTCTTCCGATCTTACCAGGGTATCTAATCCT) using Phanta Max Super-Fidelity DNA Polymerase (Vazyme, PRC) (95°C for 3 min and then 35 cycles at 95°C for 15 sec, 65°C for 15 sec, and 72°C for 1 min before a final step at 72°C for 5 min). The purified amplicons were sequenced using Miseq sequencing technology (Illumina) at the GeT-PLaGe platform (Toulouse, France). Human and mouse sequences are available under https://doi.org/10.57745/FSQYPA and https://doi.org/10.57745/8YQANH respectively. Paired-end reads obtained from MiSeq sequencing were then dereplicated and processed using the Galaxy supported FROGS version 4.1 [52]. Amplicon Sequence Variant (ASV) were clustered using Swarm with parameter d = 1 and chimeras were filtered following FROGS version 4.1 guidelines. Assignation was performed using 16S_REFseq_Bacteria_20230726 database. ASV with abundances lower than 0.0005% of the total read set were removed prior to analysis. Subsequent analysis were done using the phyloseq R packages [53]. Samples were rarefied to even sampling depths before computing within-samples compositional diversities (Chao1 and Shannon index) and between-samples compositional diversity (Bray-Curtis) Raw, unrarefied ASV counts were used to produce relative abundance graphs and to find family and genera with significantly different abundances among mouse groups or between normo-fructose absorber and fructose malabsorber individuals respectively.

##### Statistical analysis

For human study, demographics and clinical characteristics parameters were summarized as median with interquartile range (IQR) and compared between groups using the Mann– Whitney nonparametric test. All statistical analysis were performed using SAS, version 9.4 (SAS Institute, Cary, North Carolina, USA). Sugar intake data were expressed as mean ± SEM and compared between groups using the Mann–Whitney nonparametric test in GraphPad Prism 10 (San Diego, CA, USA).

For mice study, statistical analysis was performed using GraphPad Prism 10 (San Diego, CA, USA). Data are expressed as mean ± SEM and individual values are plotted on graph when possible. After normal Gaussian distribution was verified using the Kolmogorov-Smirnov test, appropriate parametric or non-parametric tests were chosen as mentioned in figure legends. For microbiota analysis: Alpha diversity index (Observed species, Chao1 and Shannon) was analyzed using 1-way ANOVA. A permutational multivariate analysis of variance (PERMANOVA) test was performed on the Bray–Curtis matrices using 9999 random permutations and at a significance level of 0.05. Phylum and family relative abundances were compared using a Mann-Whitney test. The Spearman’s rank correlation coefficients between the relative abundance of gut microbiome at family level and LPS plasma concentration, STAI score as well as fructose intake profile were determined using R software for correlational statistical analysis. All statistical tests used two-sided 0.05 significance threshold.

## RESULTS

### Volunteers with fructose-malabsorption display traits of anxiety and increased level of circulating LPS despite similar fructose intake

A total of 55 healthy volunteers were included in the study. One volunteer left the study, and five were excluded due to exclusion criteria (Supp Table S1). Sixty percent of evaluated volunteers were tested positive for fructose malabsorption (29 fructose malabsorbers (Mal-Abs) and 20 normoabsorbers (Normo-Abs)). Age, weight, and BMI did not differ between the groups (Table 2). Among them, 48 volunteers completed the dietary surveys. Food diaries did not reveal any difference in total daily fructose intake between Normo-Abs and Mal-Abs individuals, with both groups consuming on average 28.6 ± 3.4 and 32.8 ± 3.0 grams of fructose per day, respectively (Figure 1A). Fructose intake from beverages and sweet products was similar between Normo-Abs and Mal-Abs volunteers, accounting for 29% and 46% of the total fructose daily intake, respectively. Fructose intake from fruits and vegetable was also similar between Normo-Abs and Mal-Abs groups, representing about 18 % of the total fructose intake (4.4 ± 4.8 and 5.8 ± 6.6 g of fructose/day for Normo-Abs and Mal-Abs, respectively) (Figure 1A). Approximately 6 % of the fructose consumed daily originates from either cereals or dairy in both Normo-Abs and Mal-Abs individuals (3.0 ± 2.0 and 3.3 ± 1.8 g of fructose/day for Normo-Abs and Mal-Abs respectively). However, these averages concealed a wide disparity in the total amount of fructose consumed (ranging from 10 to 75 g of fructose/day for Mal-Abs and from 8 to 60 g of fructose/day for Normo-Abs) and in the profile of dietary fructose sources within the cohort, irrespective of fructose malabsorption status (Figure 1B). Notably, 14% of individuals had a daily total fructose consumption of more than 50 g/day, and 36% consumed more than 25g of added fructose daily (Supp. Figure S1). Among them, 45% of Mal-Abs individuals consumed more than 25 g of added fructose, compared to only 26% of Normo-Abs individuals.

**Figure 1:**
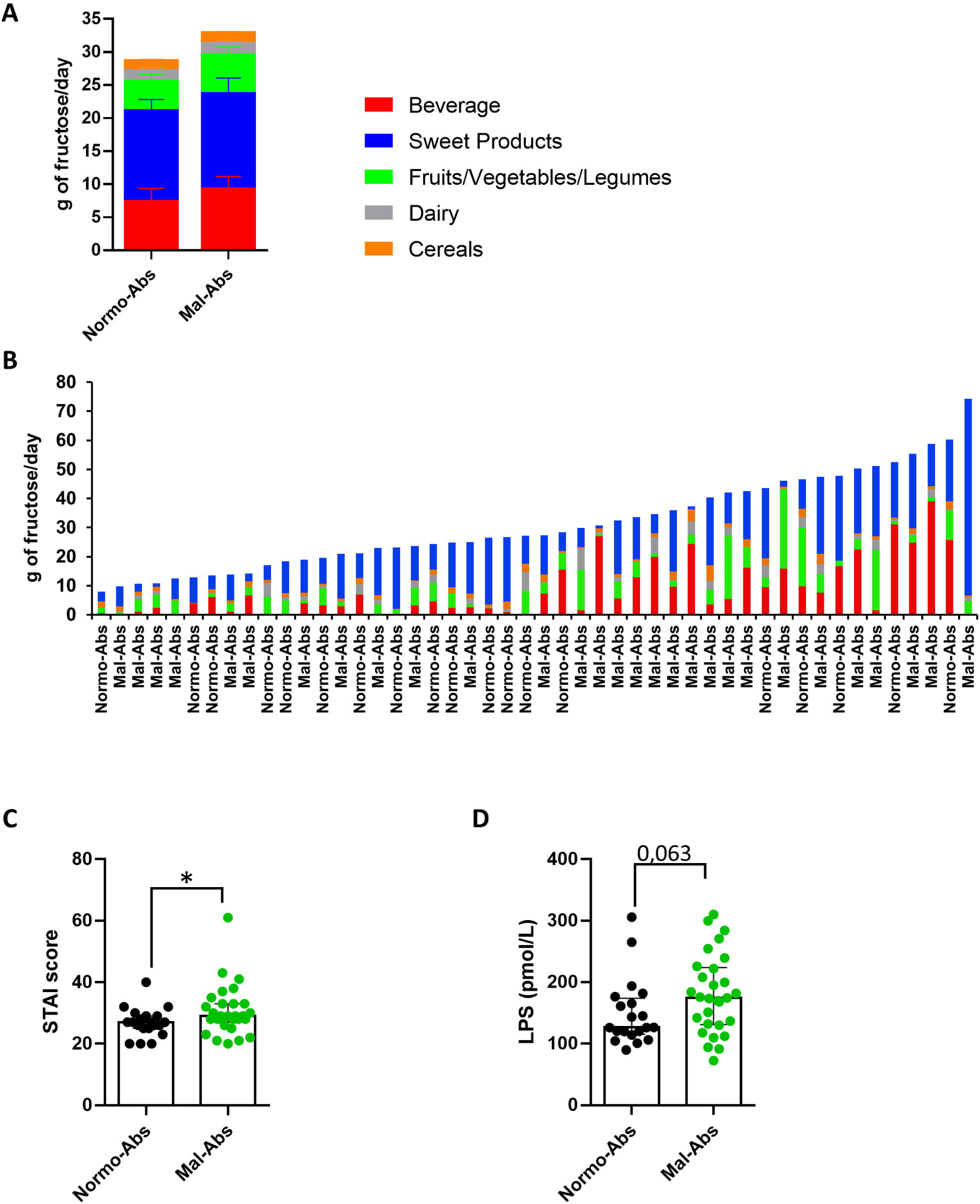
Dietary and clinical parameters of Normo-Abs and Mab-Abs volunteers. **(A)** Average daily contribution in gram of different food and beverage groups to total fructose intake in Normo-Abs and Mal-Abs volunteers. **(B)** Individual daily contribution in gram of different food and beverage groups to total fructose intake. **(C)** STAI scores and **(D)** LPS circulating levels in in Normo-Abs and Mal-Abs volunteers. Fructose intakes are means ± SEM. STAI scores and LPS levels values are median with interquartile. Fructose intakes, STAI scores and LPS levels were compared using Mann-Whitney comparison: * p < 0.05. STAI: State-Trait Anxiety Inventory, LPS: Lipopolysaccharide.

**Table 2:** Demographics and clinical characteristics.

|  | **Volunteers without**<br>**fructose**<br>**malabsorption**<br>**(Normo-Abs)** | **Volunteers with**<br>**fructose**<br>**malabsorption**<br>**(Mal-Abs)** | **p values** |
| --- | --- | --- | --- |
| **Number of volunteers** | 20 | 29 |  |
| **Age (years)** | 24.8 [21.9 ; 31.6] | 24.7 [20.2 ; 29.4] | 0.746 |
| **Weight (kg)** | 74 [71.5 ; 81.5] | 72 [65 ; 77] | 0.101 |
| **BMI (kg/m2)** | 23.7 [22.1 ; 24.3] | 22.3 [21.1 ; 24.1] | 0.108 |
| **HADS anxiety score visit1** | 5 [4 ; 6.5] | 6 [5 ; 6] | 0.586 |
| **HADS depression score visit1** | 2 [1 ; 3] | 2 [1 ; 4] | 0.337 |
| **HADS anxiety score visit2** | 4.5 [3 ; 6.5] | 5 [4 ; 7] | 0.659 |
| **HADS depression score visit2** | 2.5 [0.5 ; 4.5] | 2 [1 ; 4] | 0.702 |
| **STAI score visit2** | 25.5 [23 ; 30] | 27 [25 ; 34] | 0.059 |
| **Fecal calprotectin (µg/g)** | 27 [18.5 ; 68] | 29 [15 ; 53] | 0.589 |
| **C-reactive protein (mg/L)** | 0.4 [0.2 ; 1] | 0.5 [0.3 ; 1] | 0.474 |

Values are expressed as the median with interquartile range. Comparison between the two groups of volunteers was performed using a Mann-Whitney test. For all the parameters Normo-Abs n=20 and Mal-Abs n=29 excepted for fecal calprotectin Normo-Abs n=20 and Mal-Abs n=28

To compare anxiety traits (measured by the STAI questionnaire) in healthy volunteers with and without fructose malabsorption, we utilized the State-Trait Anxiety Inventory. At the first visit, the Mal-Abs groups exhibited a significantly higher median STAI score compared to the Normo-Abs group: 29 (IQR 28 – 33) versus 27 (IQR : 25 – 29), respectively (p=0.042) (Figure 1C). Similarly, a strong trend was observed during the second visit, despite the participants’ familiarity with the questionnaire, with the Mal-Abs group still showing higher anxiety scores (p = 0.054) (Table 2). These results indicate an increase in anxiety traits in individuals with fructose malabsorption. As shown in Table 2, HAD-depression and HAD-anxiety average scores did not significantly differ between Normo-Abs and Mal-Abs individuals and remained below 11 indicating an absence of severe depressive or anxiety conditions in the cohort. Among the circulating inflammatory parameters measured, plasma LPS levels tended to be higher in Mal-Abs compared to Normo-Abs, showing a near-significant difference (p = 0.063; Figure 1D), while fecal calprotectin and the CRP levels were similar between the two groups (Table 2), indicating no gastrointestinal or significant systemic inflammation associated with fructose malabsorption.

### Bacteria associated with fructose malabsorption and traits of anxiety

Changes in gut microbiota composition can modulate the levels of circulating LPS [54]. We therefore analyzed the fecal microbiota composition of Normo-Abs and Mal-Abs volunteers. For a taxonomic overview, Figure 2A shows the relative abundance of bacterial taxa aggregated at the family levels. Taxonomic assignment at family (Figure 2A) or phylum level (Supp. Figure S2A) revealed no difference in bacterial communities between Normo-Abs and Mal-Abs groups. We found that the fecal microbiome was dominated by the family *Lachnospiraceae* (∼39% in both groups), *Oscillospiraceae* (20.5% and 18.5% in Normo-Abs and Mal-Abs, respectively) and *Bifidobacteriaceae* (9.7% and 14.3% in Normo-Abs and Mal-Abs, respectively). *Lachnospiraceae* and *Oscillospiraceae* both belong to the Bacillota phylum, which represents more than 60% of the total microbiome composition. Fecal microbiota β-diversity composition did not differ significantly between Normo-Abs and Mal-Abs volunteers (Figure 2B). Similarly, the community richness (Chao1 index) and diversity (Shannon index) remained unchanged between the two groups (Figure 2C). However, at the genus level, 2 taxa, *Agathobacter* and *Bifidobacterium* displayed a significant higher relative abundance in Mal-Abs individuals, while the three genera *Prevotella*, *Enterococcus* and *Zhenpiania* were significantly decreased (Figure 2D).

**Figure 2:**
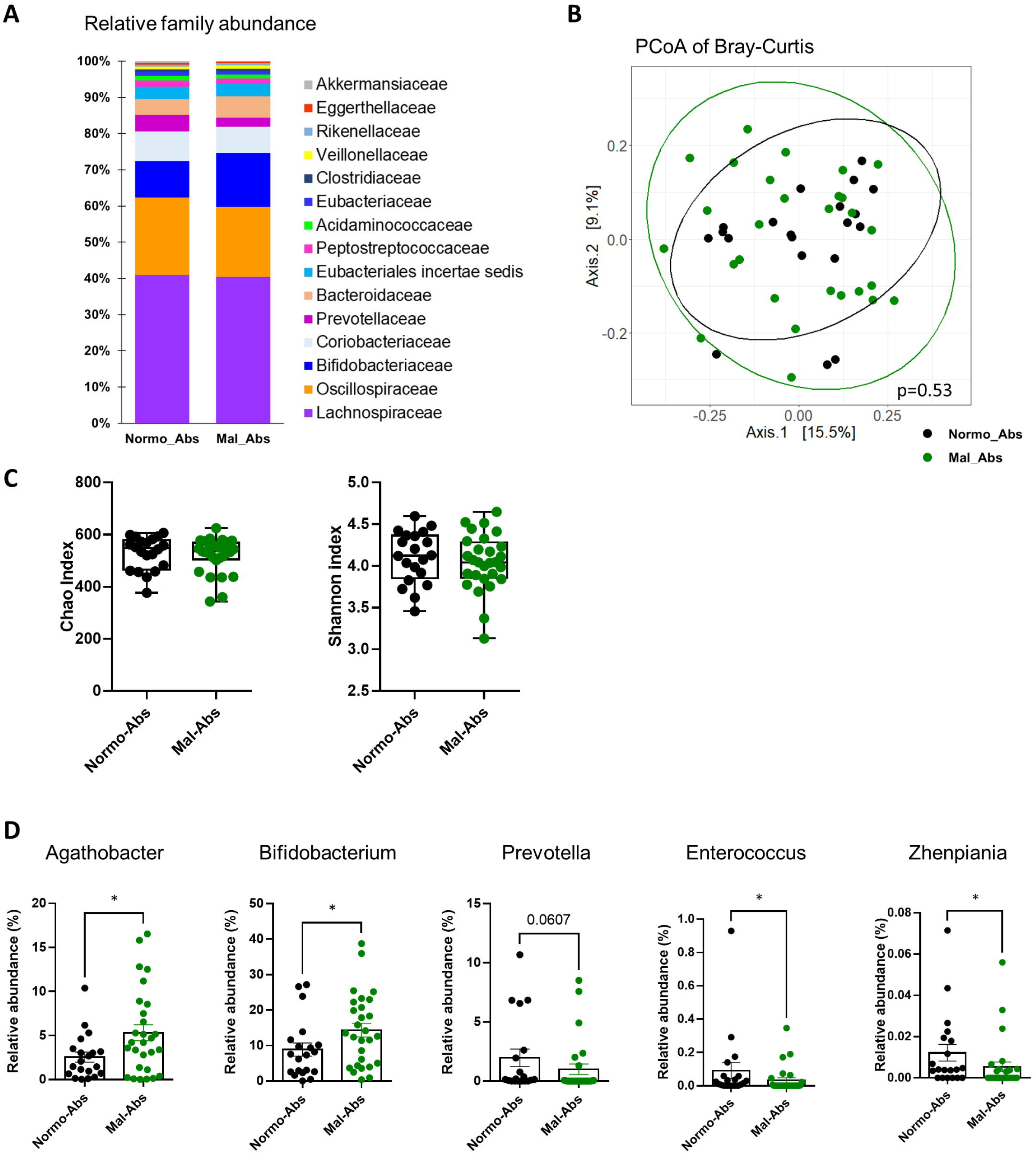
Microbiota composition of fecal samples of Mal-Abs and Normo-Abs volunteers. **(A)** Taxonomic overview of the fecal microbiome at the family level for unrarefied data. Bar plots display the relative abundance of the top 15 taxa with the highest average abundance across group average. **(B)** Principal coordinates analysis (PCoA) of Bray-Curtis compositional dissimilarity. Each dot represents of one volunteer. **(C)** Chao1 and Shannon Index as indicators of a-diversity. **(D)** The differential genus level abundances of the fecal microbiota. Chao1 index, Shannon Index and relative genera abundance values are means ± SEM. Genera abundance data were compared using Mann-Whitney comparison: * p < 0.05.

Interestingly, the abundance of certain bacterial taxa correlated with fructose consumption from specific sources, independent of fructose malabsorption status (Figure 3A). For example, *Streptococcaceae* was positively correlated with fructose intake from dairy products, supporting the accuracy of food questionnaire responses, as species from this family have previously been linked to high-dairy diets [55]. *Coprobacillaceae* showed a positive correlation with fructose intake from beverages. Additionally, a cluster of four families exhibited positive correlations with different types of added fructose intake: *Barnesiellaceae* and *Peptococcaceae* correlated with fructose from sweet beverages and total added fructose; *Sphingobacteriaceae* correlated with total, added, and beverage-derived fructose; and *Erysipelotrichaceae* correlated with total fructose. In contrast, some families were negatively correlated with fructose intake—*Eubacteriaceae* with fructose from beverages and *Atopobiaceae* with total, added, and beverage-derived fructose. Fructose from fruits and vegetables was negatively correlated with *Turicibacteraceae*, which, along with *Bacteroidaceae*, showed a strong positive association with STAI scores (Figure 3B). Conversely, *Eubacteriaceae* and *Erysipelotrichaceae* were negatively correlated with anxiety traits. Finally, *Christensenellaceae*, *Gracilibacteraceae*, and *Rikenellaceae* were positively correlated with LPS plasma levels.

**Figure 3:**
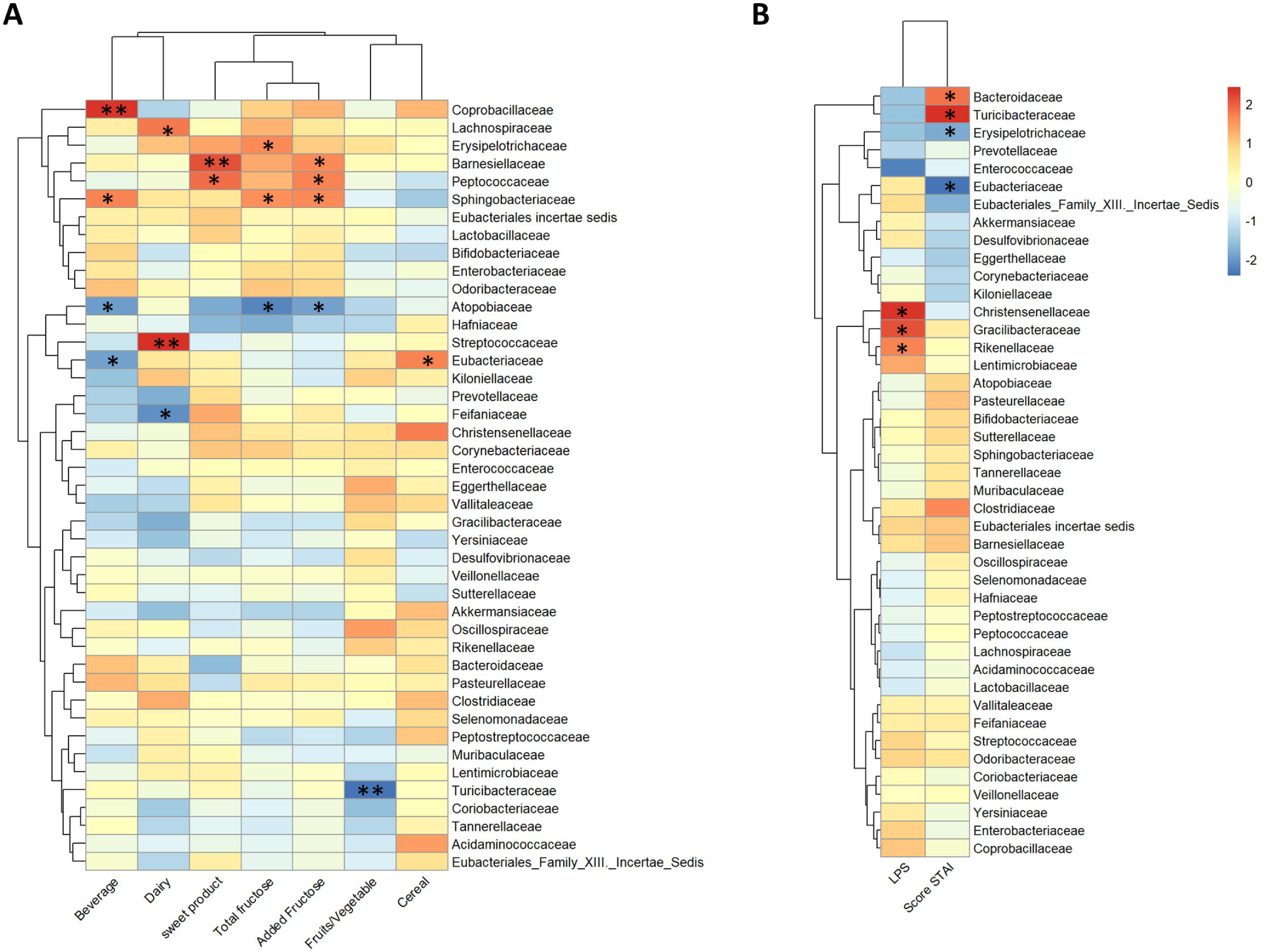
Spearman’s correlation. **(A)** Heat map of Spearman’s correlation showing significant correlations of gut microbiota at the family level and fructose intake sources. **(B)** Heat map of Spearman’s correlation showing significant correlations of gut microbiota at the family level and STAI index and LPS plasma levels. Significant correlations are shown as red (positive) or blue (negative). **\****p* < 0.05; **\*\****p* < 0.01.

### Fructose malabsorption triggers anxiety-like behaviors and alters microglia function in mice

The impact of fructose malabsorption on emotional behaviors was assessed in malabsorptive GLUT5-KO mice. Malabsorptive GLUT5-KO mice (GLUT5KO_5F) were fed a 5% fructose diet for 4 weeks before assessing anxiety- and depressive-like behaviors. WT mice can fully absorb up to 20% of dietary fructose, and previous studies have shown that microbiota composition remains unchanged when fructose intake varies between 0% and 20%, provided that only the fructose amount is modified [15]. We confirmed these findings, as WT mice fed 0% or 5% fructose for 10 days exhibited no significant changes in fecal microbiota composition at the phylum or family levels (Supp. Figure S3A). Additionally, neither WT nor GLUT5KO mice displayed differences in body weight or food intake between the 0% and 5% fructose diets, ruling out potential metabolic and behavioral confounding effects (Supp. Figure S3B and S3C). Therefore, WT mice fed 5% fructose (WT5F) as well as GLUT5-KO mice fed 0% fructose (GLUT5KO_0F) served both as control groups without fructose malabsorption. In the EPM test, GLUT5KO_5F mice exhibited fewer entries, as well as reduced distance traveled and time spent in the aversive open arm compared to normo-fructose-absorbing WT5F controls (Figure 4A), with no changes in total distance traveled during the test (Supp. Figure S4A). GLUT5KO_5F mice exhibited increased immobility time during the FST (Figure 4B). All together, these data suggest that fructose malabsorption increases anxiety- and depressive-like behaviors in Mice.

**Figure 4:**
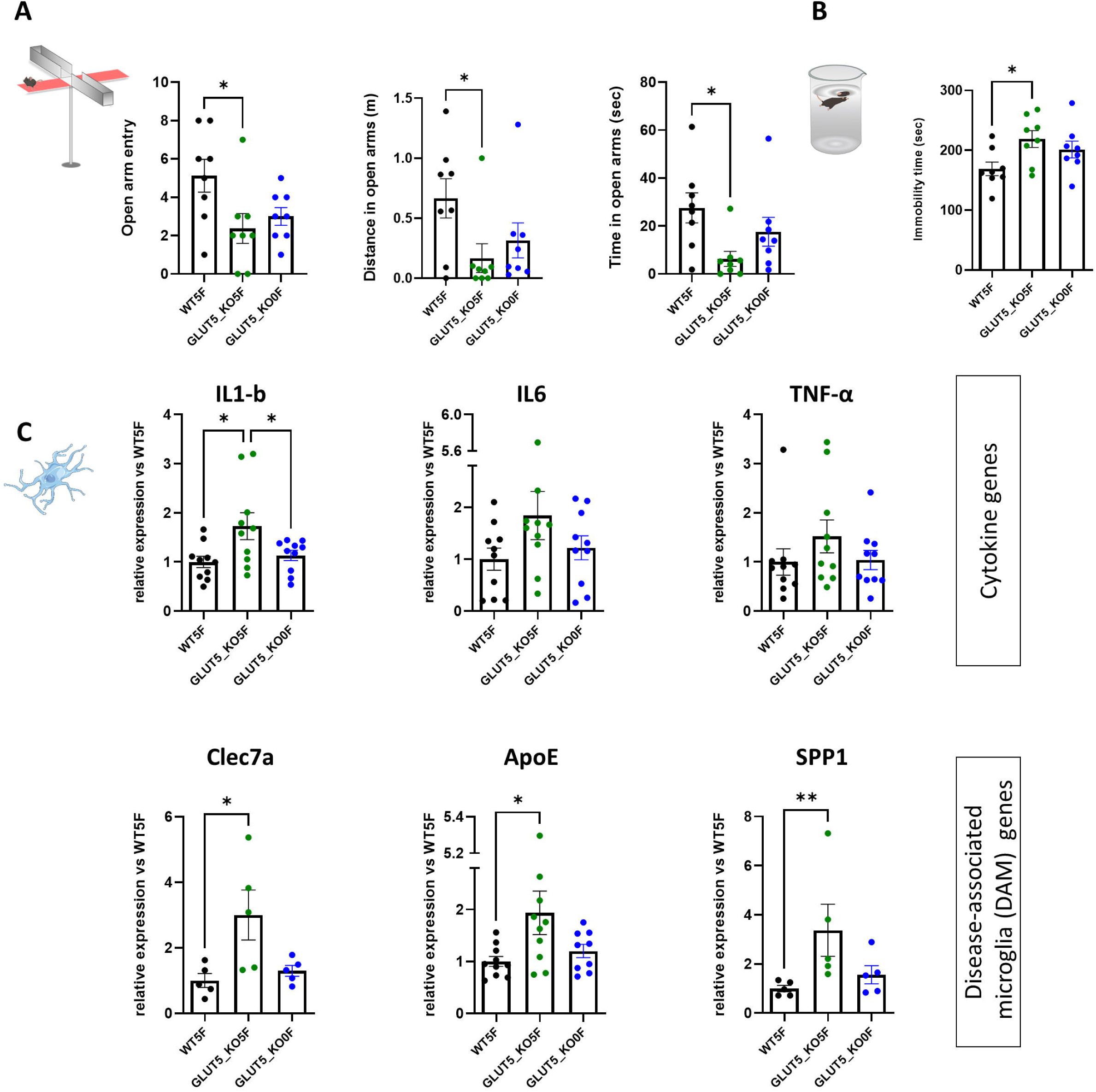
Fructose mal-absorption increases anxiety- and depressive-like behaviors and impairs microglia physiology. **(A, B)** Number of entries (A, left panel), time spent (A, middle panel) and distance traveled (A, right panel) in the open arm in the EPM and immobility time during the FST (B) of WT (WT5F, n = 8) or GLUT5_KO mice (GLUT5_KO5F, n = 8) fed a fructose 5% diet for 4 weeks or GLUT5_KO mice (GLUT5_KO0F, n = 8) fed a control diet without any fructose for 4 weeks. **(C)** Expression of cytokines (IL1-b, IL6, TNF-α) or disease-associated microglia (DAM) genes (Clec7a, ApoE, SPP1), normalized to b2m gene expression quantified in microglia isolated from brains of WT (WT5F, n = 5 to 10) or GLUT5_KO mice (GLUT5_KO5F, n = 5 to 10) fed a fructose 5% diet for 4 weeks or GLUT5_KO mice (GLUT5_KO0F, n = 5 to 10) fed a control diet without any fructose for 4 weeks. * p < 0.05; ** p < 0.01: One-way ANOVA followed by Holm-Sidak’s multiple comparisons test.

To investigate whether these behavioral changes were linked to microglia-associated neuroinflammation, we assessed the expression of selected genes in isolated microglia from normo- and malabsorptive fructose-fed mice. Expression of interleukin 1 β (Il1β) was significantly elevated in microglia from GLUT5KO_5F mice compared to the two control groups (WT5F and GLUT5KO_0F) (Figure 4C, upper panel). Additionally, Il6 expression showed an increasing trend in GLUT5KO_5F microglia relative to WT5F (p = 0.067; Figure 4C, upper panel). Fructose malabsorption also influenced the expression of disease-associated microglia (DAM) signature genes. Clec7a, Apolipoprotein E (ApoE), and osteopontin/secreted phosphoprotein 1 (Spp1) were significantly upregulated in GLUT5KO_5F microglia compared to WT5F (Figure 4C, lower panels), suggesting that fructose malabsorption modulates microglial phenotypes and functions. Furthermore, Csf1r expression showed a decreasing trend after fructose exposure in GLUT5KO_5F relative to WT5F (p = 0.054), while homeostatic genes P2ry12 and Tgfbr2 remained unchanged (Supp. Figure S4B). Interestingly, Csf1r and P2ry12 expression were both reduced in GLUT5KO_5F microglia compared to GLUT5KO_0F (Supp. Figure S4B), suggesting a GLUT5-independent mechanism underlying microglial responses to fructose feeding.

### Fructose malabsorption is associated with intestinal dysbiosis in mice

Microbiota-derived metabolites can modulate microglia functions and neuroinflammation although the exact mechanism remains elusive [17, 23]. We investigated the response of the cecal bacterial ecosystem to fructose malabsorption which parallels the alteration of mood disorder and neuroinflammation observed in GLUT5_KO5F mice. Bray-Curtis dissimilarity matrices showed that fructose malabsorption contributed significantly (p < 0.01) to the different compositions observed in microbial communities of the caecum in GLUT5_KO5F compared to WT5F and GLUT5_KO0F mice (Figure 5A). In addition, the community richness (Chao1 index) and diversity (Shannon index) was significantly increased between WT5F and GLUT5_0F but remain unchanged GLUT5_0F and GLUT5_5F indicating that fructose malabsorption did not affect the alpha-diversity of the cecal bacterial ecosystem (Figure 5B). Taxonomic assignments at family (Figure 5C) or phylum (Supp. Figure S2B) level revealed differences in bacterial communities in mice malabsorbing fructose (GLUT5KO_5F) compared to the 2 other groups. At the phylum level, the relative abundance of Actinomycetota and Bacteroidota was significantly higher in the cecum of GLUT5KO_5F mice compared to WT5F or GLUT5KO_0F mice, respectively (Supp. Figure S2B). Significant changes in relative abundance of ten families were specifically observed in the caecum of GLUT5KO_5F mice when compared to both controls WT5F and GLUT5KO_0F mice (Figure 5D and Supp. Table S2). *Peptostreptococcaceae* belonging to the Bacillota phylum (formerly known as Firmicutes) almost entirely disappeared in response to fructose malabsorption. *Streptococcaceae* and *Christensenellaceae* (belonging to the Bacillota phylum) both described as health-beneficial bacteria displayed a 10-fold relative abundance decrease in response to fructose malabsorption. By opposition, in the same phylum, *Eubacteriales_Family_XIII._Incertae_Sedis* and *Erysipelotrichaceae* relative abundance increased GLUT5KO_5F mice when compared to WT5F and GLUT5KO_0F mice. In addition, *Prevotellaceae* (Bacteroidota phylum) relative abundance increased ∼6 times more and became the fourth most abundant family in the caecum of the GLUT5KO_5F. In the same phylum the *Rikenellaceae* relative abundance was divided by two and three in the caecum of the GLUT5KO_5F mice when compared to WT5F and GLUT5KO_0F mice respectively. Finally, the *Corynebacteriaceae* (Actinomycetota phylum formerly known as Actinobacteria) increased by about 2 times in mal-absorptive GLUT5KO_5F as compared to WT5F and GLUT5KO_0F mice.

**Figure 5:**
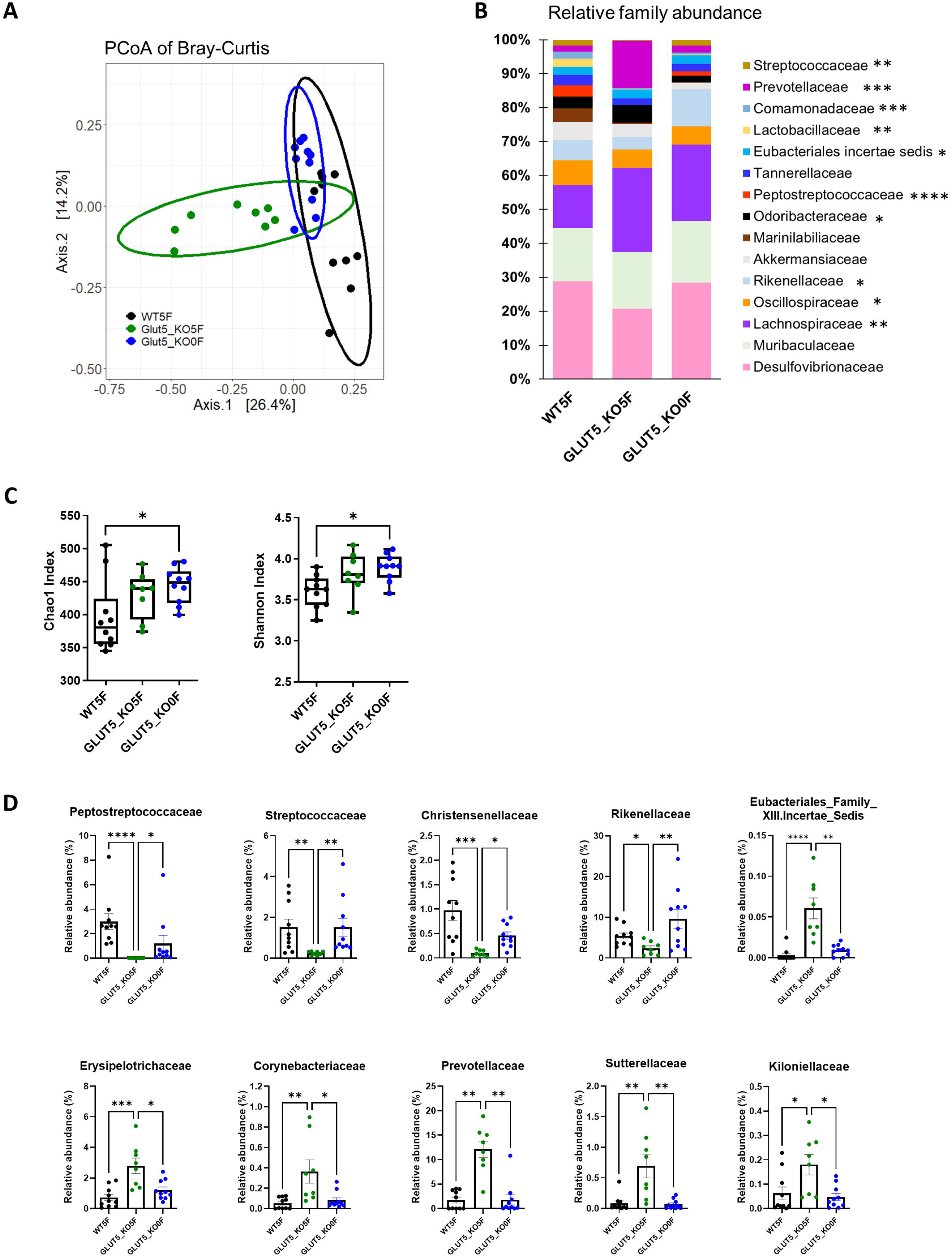
Microbiota composition of cecal sample of WT and GLUT5_KO mice. **(A)** Principal coordinates analysis (PCoA) of Bray-Curtis compositional dissimilarity. Each dot represents one mouse **(B)** Taxonomic overview of the fecal microbiome at the family level for unrarefied data. Bar plots display the relative abundance of the top 15 taxa with the highest average abundance across group average. **(C)** Chao1 and Shannon Index as indicators of a-diversity. **(D)** Relative abundances of taxa at the family level that significantly differed in GLUT5_KO5F groups. Chao1 index, Shannon Index and relative family abundance values are means ± SEM with n=8 to 10. Chao1 and Shannon Index means were compared by one-way ANOVA followed by THolm-Sidak’s multiple comparisons test. For figure B family abundance data were compared using Kruskal-Wallis test with p * < 0.05, ** p < 0.01, *** p < 0.001 and **** p < 0.0001 (multiple test detail are presented in TableS2). For Figures D family abundance data were compared using Kruskal-Wallis test followed by Dunn’s multiple comparison test: p * < 0.05, ** p < 0.01, *** p < 0.001 and **** p < 0.0001.

## DISCUSSION

Clinical and pre-clinical data from this study show that fructose malabsorption is associated with alterations in gut microbiota and increased anxiety levels. Using animal models, we also show that these impairments associated with fructose malabsorption correlate to changes in microglia gene expression related to neuroinflammation. Our study further reveals that more than 50% of healthy individuals in our cohort malabsorb fructose, regardless of their intake of fructose-containing foods. Additionally, about 35% of them exceeded the recommended limits for fructose consumption.

### Fructose Intake Patterns in Normo- and Malabsorbers

While most countries report total sugar intake, fewer provide specific data on added sugars and sucrose, and even fewer detail the precise intake of fructose [35, 56, 57]. Although recent efforts have combined food databases with composition data to assess specific sugar consumption [8], a significant gap remains. Our study helps addressing this gap by providing comprehensive data on total fructose intake and its dietary sources in humans. In the MoodyFructose cohort, we report an average daily fructose intake of 30 g per individual. These data are coherent with previous European or French surveys for this age range, even when the total intake of fructose was extrapolated from the total intake of sucrose [57]. Despite the cohort’s homogeneity in terms of age, sex, and BMI, we observed significant heterogeneity in the food sources of fructose-containing sugars among individuals, providing valuable insights into fructose intake patterns and their potential health implications. In 2015, the World Health Organization assessed the effects of dietary sugars on health and recommended that free sugar intake should be less than 10%, or even 5%, of total energy for both children and adults, which represents about 50 grams (or even 25 grams for the lower recommended limit) of free sugar per day [58]. Since fructose accounts for approximately 50% of total sugar intake, these recommendations translate to a maximum recommended added fructose intake of 25g per day. Based on these recommendations, nearly 40% of individuals in the MoodyFructose cohort exceed this upper limit. Additionally, in 2018, the ANSES institute set a maximum limit for fructose intake at 50 g/day [7], and notably, 14% of individuals in our cohort surpassed this threshold in their daily total fructose consumption.

### Fructose malabsorption and anxiety

Although fructose overconsumption is recognized by health institutes as potentially harmful, these recommendations mainly focus on its effects on energy metabolism or peripheral tissues [7, 8, 58]. There is no human data indicating that fructose overconsumption impairs brain function or health. However, numerous animal studies have shown that fructose-enriched diets lead to impaired brain health and increased anxiety or depressive-like behaviors [30, 59, 60]. In some cases, these brain dysfunctions are linked to increased neuroinflammation [29, 30, 60]. Fructose may affect the brain by promoting neuroinflammation through pathways like direct effects on brain cells, such as microglia (where the fructose transporter Glut5 is highly expressed) [61], or by influencing the gut microbiome, as shown in rodent studies [15, 29, 62–68]. The current study examined the impact of fructose malabsorption in human and animal models to test the hypothesis that fructose malabsorption-induced gut dysbiosis contributes to increased neuroinflammation and impaired emotional behaviors. We demonstrated that, regardless of overall fructose intake volunteer with fructose malabsorption or GLUT5_KO mice fed a moderate amount of fructose exhibit increased anxiety symptoms. Although most volunteers in the Moodyfructose cohort had STAI scores between 20 and 37 (indicating ‘no or low anxiety’) [41], those with fructose malabsorption exhibited a slight but significant increase in anxiety state compared to normo-absorbers. These findings suggest that fructose malabsorption is associated with anxiety traits, without reaching pathological levels, as supported by the low HAD scores.

### Role of gut microbiota in fructose malabsorption-associated anxiety

The link between gut microbiota and emotional disorders has been suggested with studies showing significant differences in fecal microbiota between depressed patients and healthy individuals [21, 69–73]. Rodent studies further support this connection, as microbiota transfer from depressed patients induces mood disturbances in recipient mice [70, 73]. However, no definitive microbiome signature for anxiety or depression has been identified [71]. Moreover, most human studies focus on major anxiety or depressive disorders [71], whereas the present study explores the potential link between fructose malabsorption- associated microbiota and anxiety traits. We found significant differences in five bacterial genera between fructose malabsorbers and normoabsorbers, with Bifidobacterium among the most affected taxa. Oligo-, di-, and monosaccharides promote the growth of *Bifidobacterium* [74], and its increase in mouse models of moderate fructose malabsorption suggests it may serve as a microbial marker [15]. While *Bifidobacterium* is often considered beneficial for anxiety and depression in both rodents and humans [75], another study reported a negative correlation with mood in humans [76]. Additionally, *Prevotella*, which was lower in fructose malabsorbers, has been linked to positive mood [76]. A reduction in *Prevotella* has also been associated with Westernized diets (characterized by a reduced amount of microbiota-accessible carbohydrate) and the loss of carbohydrate-active enzymes, suggesting that fructose malabsorbers may have a reduced capacity to metabolize certain carbohydrates [77]. In contrast, *Prevotella* was drastically increased in GLUT5-KO mice fed 5% fructose diet, where excess fructose reached the lower intestine and likely became a microbiota-accessible carbohydrate [78]. Indeed, unlike human malabsorbers who did not consume more fructose than normoabsorbers, Glut5-KO mice were fed a 5% fructose diet. We also found bacterial families positively (*Turicibacteraceae* and *Bacteroidaceae*) or negatively (*Eubacteriaceae* and *Erysipelotrichaceae*) associated with anxiety traits. *Bacteroidaceae*, linked to anxiety in IBS patients, is lower in major depression disorder [79, 80], while *Turicibacter* correlates with cognitive impairment in both humans and mice [81, 82]. Interestingly, fructose intake from fruits/vegetables negatively correlated with *Turicibacteraceae* abundance. Similarly, *Eubacteriaceae*, negatively associated with anxiety traits, showed an inverse correlation with fructose intake from beverages. These findings highlight the importance of fructose sources in shaping gut microbiota and mood regulation, consistent with prior studies [83]. Finally, we found that fructose malabsorption tend to increased human LPS and pro-inflammatory cytokine levels in mouse microglial cells, alongside altered mood parameters. *Rikenellaceae,* a gram-negative bacterial family, is one of the taxa positively correlated with LPS levels in our cohort, and previous studies have established a link between LPS and the *Rikenellaceae* family [84]. However, in mice, *Rikenellaceae* decreased in response to fructose diet in GLUT5_KO, suggesting potential alternative mechanisms linking microbiota changes to neuroinflammation in the context of fructose malabsorption.

### Fructose malabsorption alters microglia and increases neuroinflammation

In the present study, we show changes in selected microglia gene expression in mal-absorptive Glut5-KO mice fed 5% fructose, suggesting altered microglia physiology is. We observed a significant increase in IL1β expression, along with a trend towards increased IL6 expression, indicating that fructose malabsorption induces basal neuroinflammation. Given that fructose is poorly absorbed and that fructose-fed GLUT5_KO exhibit dysbiosis, our data align with studies showing that fructose overfeeding is associated with increased cytokine expression in specific brain areas [29, 30, 60]. In addition, we found that malabsorptive GLUT5_KO5F showed increased expression of *Clec7a*, *ApoE* and *Spp1*, genes associated with DAM signature, and that are primarily linked to phagocytic phenotype and function of microglia. Alterations in microglia phagocytic functions, coupled with neuroinflammation, may impact surrounding brain cells and their activities in homeostasis and disease, including neurodegenerative diseases [85]. This aligns with studies suggesting that fructose overfeeding increases the expression of Alzheimer’s disease markers [86, 87] and that fructose consumption is associated with a higher risk of Alzheimer disease in humans [88, 89]. Further analyses are needed to better understand the impact of fructose malabsorption on neuroinflammation. Importantly, we also found an altered homeostatic signature of microglia in Glut5-KO mice, independent of fructose (Supp. Figure S3). Since GLUT5 is a homeostatic marker [61], this suggests that the deletion of Glut5 in microglia may sensitize them to acquire a DAM phenotype. Additional work using conditional deletion of *Glut5* in gut epithelial cells or microglia is necessary to further elucidate the impact of fructose malabsorption or GLUT5 itself on microglia-dependent control of neuroinflammation.

### Conclusions

In conclusion, both human and preclinical models of fructose malabsorption have demonstrated increased anxiety traits in healthy volunteers and anxiety-like behaviors in fructose malabsorptive mice, respectively. In both cases, anxiety was associated with an unbalanced intestinal bacterial ecosystem and the presence of inflammation. These findings suggest that a better understanding of the mechanistic relationship between inflammatory markers and bacterial taxa in the context of depression and anxiety could aid in the management of patients in the future. However, the current study did not explore, in human, the impact of increased fructose consumption in the context of fructose malabsorption condition, which remains an important area for further research. Specifically, whether chronic fructose malabsorption—either with or without increased fructose consumption—could exacerbate trait anxiety and contribute to the deterioration of mental health remains unclear. Investigating this question could improve nutritional recommendations for fructose consumption and offer new insights into the role of diet in mental health.

## DECLARATIONS

### ETHICS APPROVAL AND CONSENT TO PARTICIPATE

Moodyfructose human cohort was approved by the national ethical committee “Comité de protection des personnes Ouest II*”* on 25^th^ of June 2021 and was registered at www.clinicaltrials.gov (NCT05371067). All volunteers gave written informed consent before participating. The study protocol conformed to the ethical guidelines of the 1975 Declaration of Helsinki. All rodent experimental procedures were conducted in accordance with the European directive 2010/63/UE and approved by the French Ministry of Research and local ethics committees (APAFIS#: 19870 and APAFIS#23075).

### CONSENT FOR PUBLICATION

Not applicable.

### AVAILABILITY OF DATA AND MATERIALS

All data are included in this article and its supplementary information files.

### AUTHORS’ CONTRIBUTIONS

CM, XF and VD designed and supervised the project and wrote the manuscript. All authors edited and approved the manuscript. AC, DP and VD performed rodent experiments. CMa, AC performed microglia analysis. LR, LN and VD performed analysis of rodent behavior experiments. JB and JPPdB analyzed LPS. MMo and VD performed microbiota analysis. AML and MPT included volunteers. AML, GG, CG, VB and MMa acquired clinical data. MQM were responsible of the samples and storage. OG and EH performed statistics on clinical data. CM, VD, XF, AC, SL, CMa, AML, MPT, LR, LN, GG and OG edited/reviewed the manuscript.

## ACKNOWLEDGMENTS

XF and AC are grateful to Bordeaux INP and INRAE for their support. Authors are thankful to the following: (i) the CIRCE (Behavioural Engineering Centre) facility of Bordeaux Neurocampus and at the IERP platform, (INRAE, UE0907, Jouy-en-Josas, France) for their technical assistance, (ii) the RRI Food4BrainHealth for the discussions of the data, (iii) the INRAE MIGALE bioinformatics platform (http://migale.jouy.inra.fr) for providing computational resources, (iv) the staff of GeT-PLaGe platform (Toulouse, France) and (v) Céline André, Virginie Buchbach, Maxime Depierre, Nell Marty and Hélène Renaux, Odile Vandapel for their technical help during the clinical study.

