## Supplementary data for "Fructose malabsorption induces dysbiosis and increases anxiety in Human and animal models"

### Slide 1
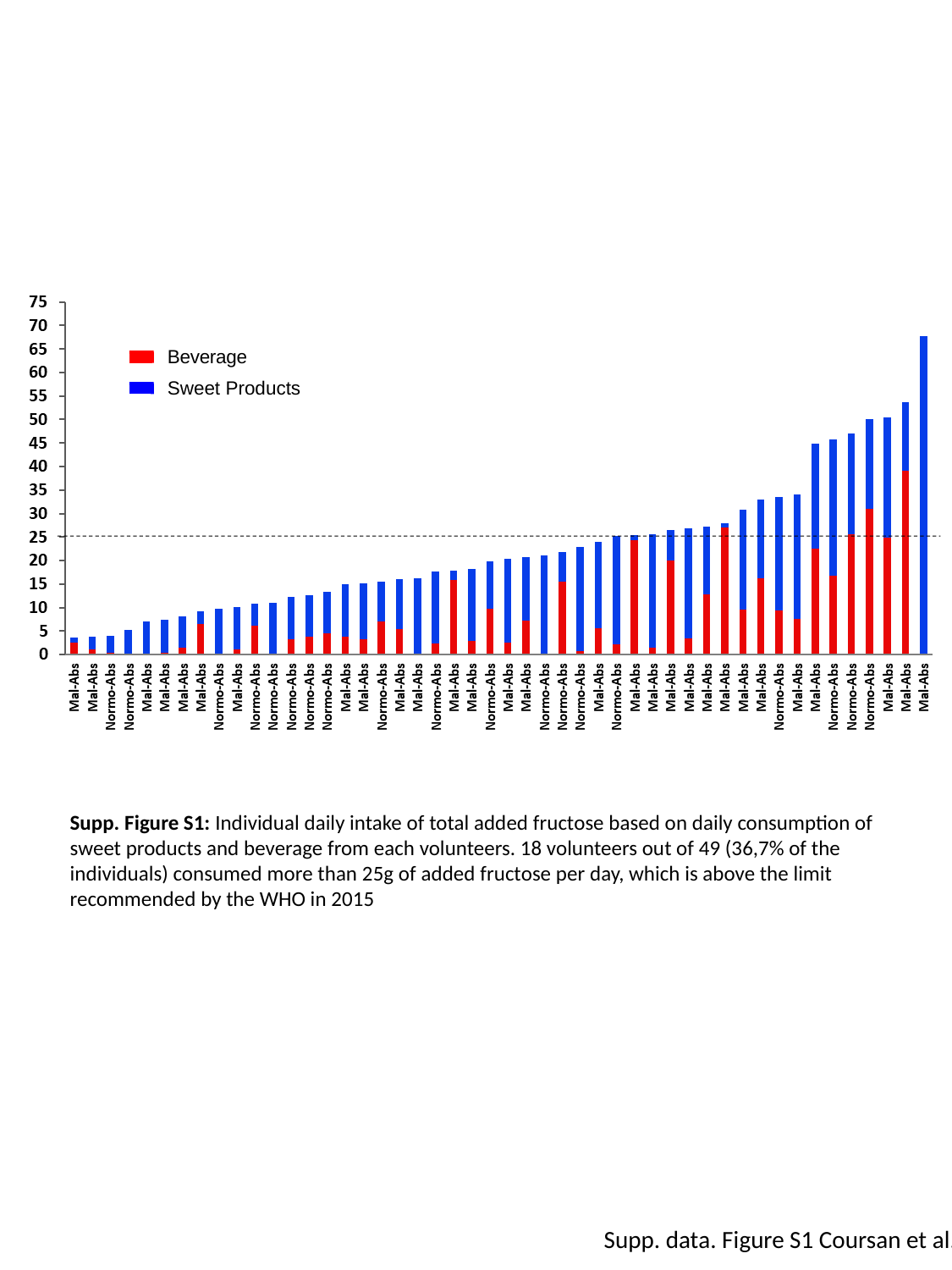

Supp. Figure S1: Individual daily intake of total added fructose based on daily consumption of sweet products and beverage from each volunteers. 18 volunteers out of 49 (36,7% of the individuals) consumed more than 25g of added fructose per day, which is above the limit recommended by the WHO in 2015
Supp. data. Figure S1 Coursan et al.

### Slide 2
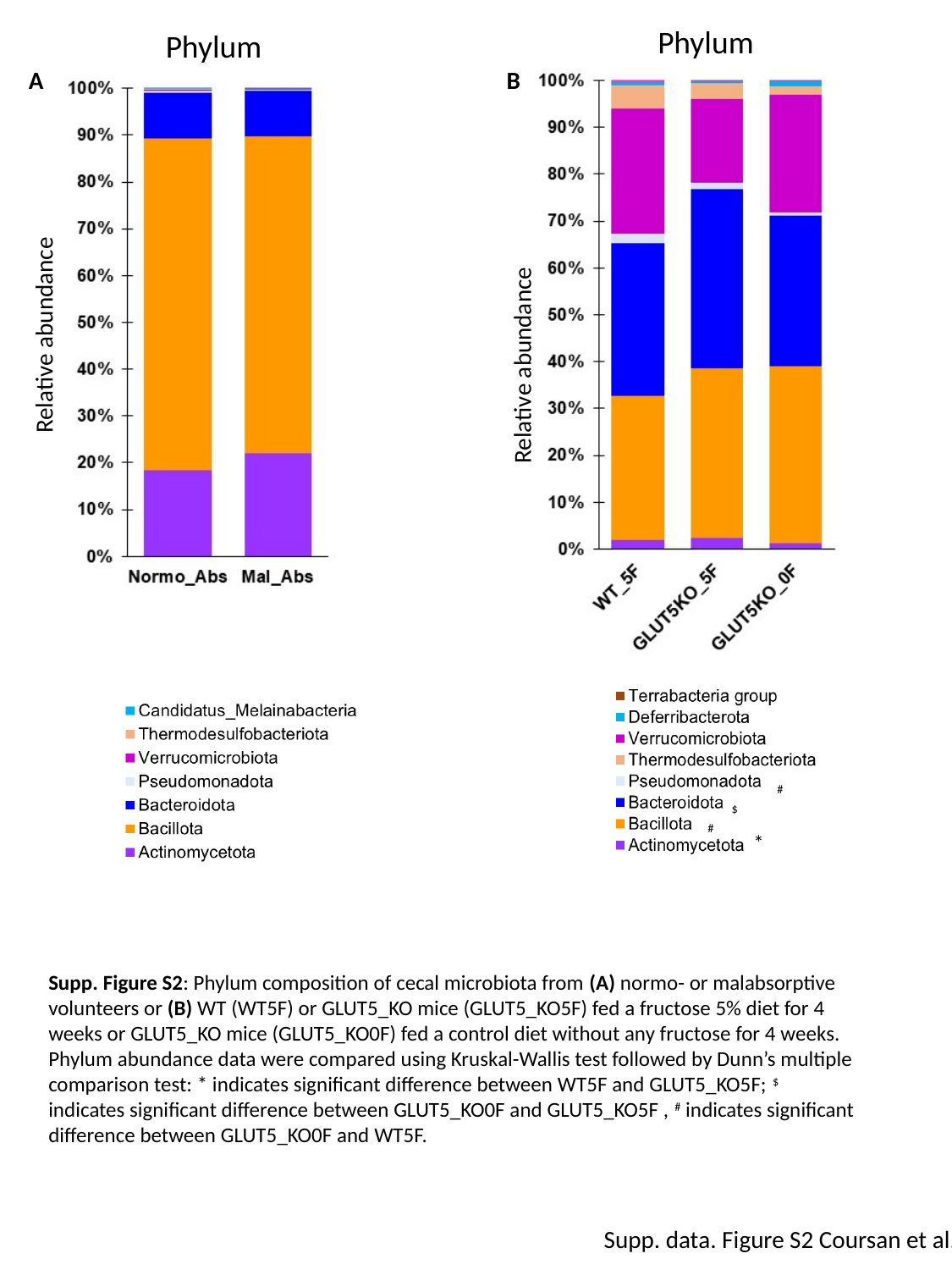

Phylum
Phylum
A
B
Relative abundance
Relative abundance
^#^
^$^
^#^
*
Supp. Figure S2: Phylum composition of cecal microbiota from (A) normo- or malabsorptive volunteers or (B) WT (WT5F) or GLUT5_KO mice (GLUT5_KO5F) fed a fructose 5% diet for 4 weeks or GLUT5_KO mice (GLUT5_KO0F) fed a control diet without any fructose for 4 weeks. Phylum abundance data were compared using Kruskal-Wallis test followed by Dunn’s multiple comparison test: * indicates significant difference between WT5F and GLUT5_KO5F; ^$^ indicates significant difference between GLUT5_KO0F and GLUT5_KO5F , ^#^ indicates significant difference between GLUT5_KO0F and WT5F.
Supp. data. Figure S2 Coursan et al.

### Slide 3
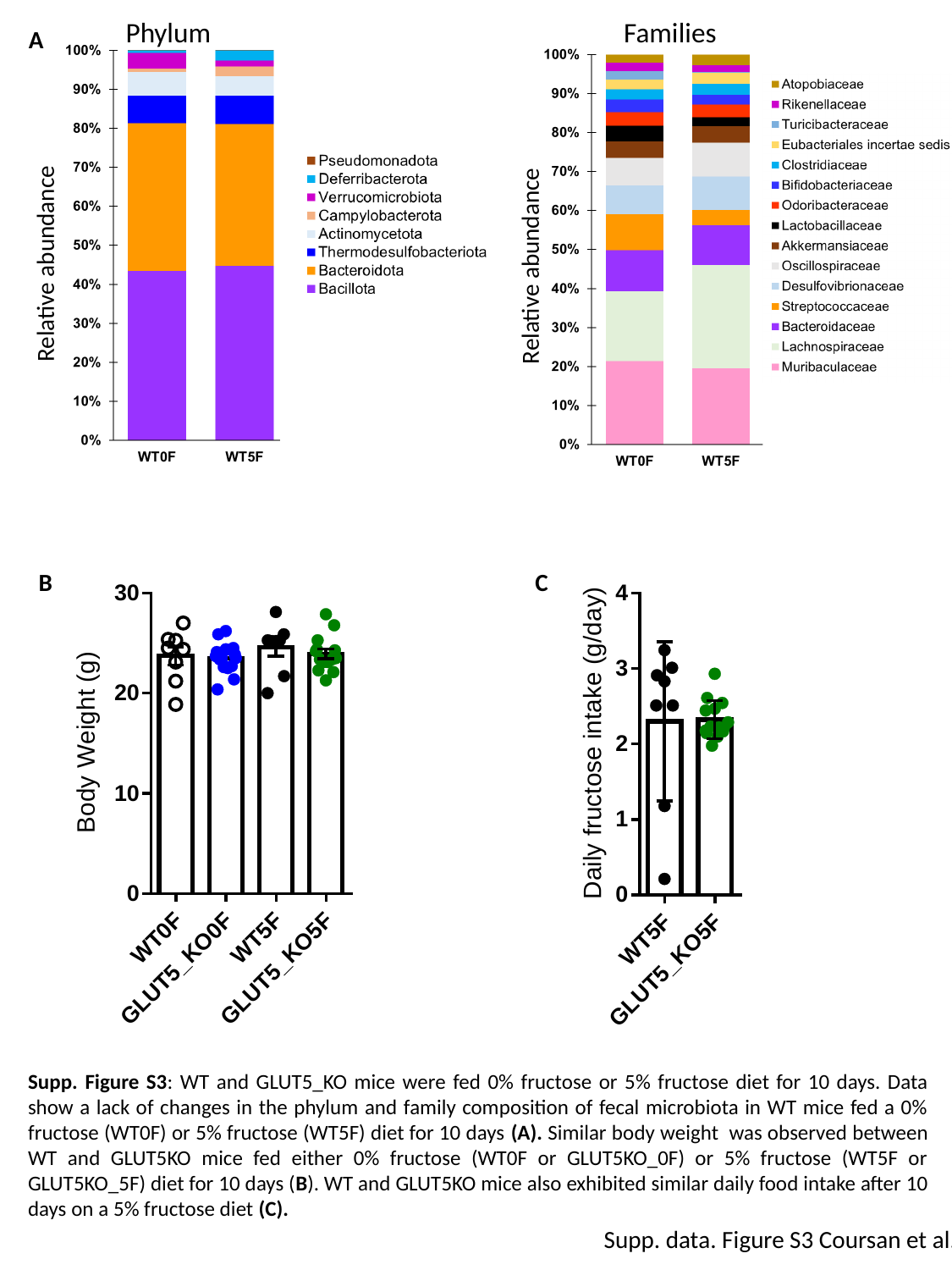

Phylum
Families
A
Relative abundance
Relative abundance
B
C
Supp. Figure S3: WT and GLUT5_KO mice were fed 0% fructose or 5% fructose diet for 10 days. Data show a lack of changes in the phylum and family composition of fecal microbiota in WT mice fed a 0% fructose (WT0F) or 5% fructose (WT5F) diet for 10 days (A). Similar body weight was observed between WT and GLUT5KO mice fed either 0% fructose (WT0F or GLUT5KO_0F) or 5% fructose (WT5F or GLUT5KO_5F) diet for 10 days (B). WT and GLUT5KO mice also exhibited similar daily food intake after 10 days on a 5% fructose diet (C).
Supp. data. Figure S3 Coursan et al.

### Slide 4
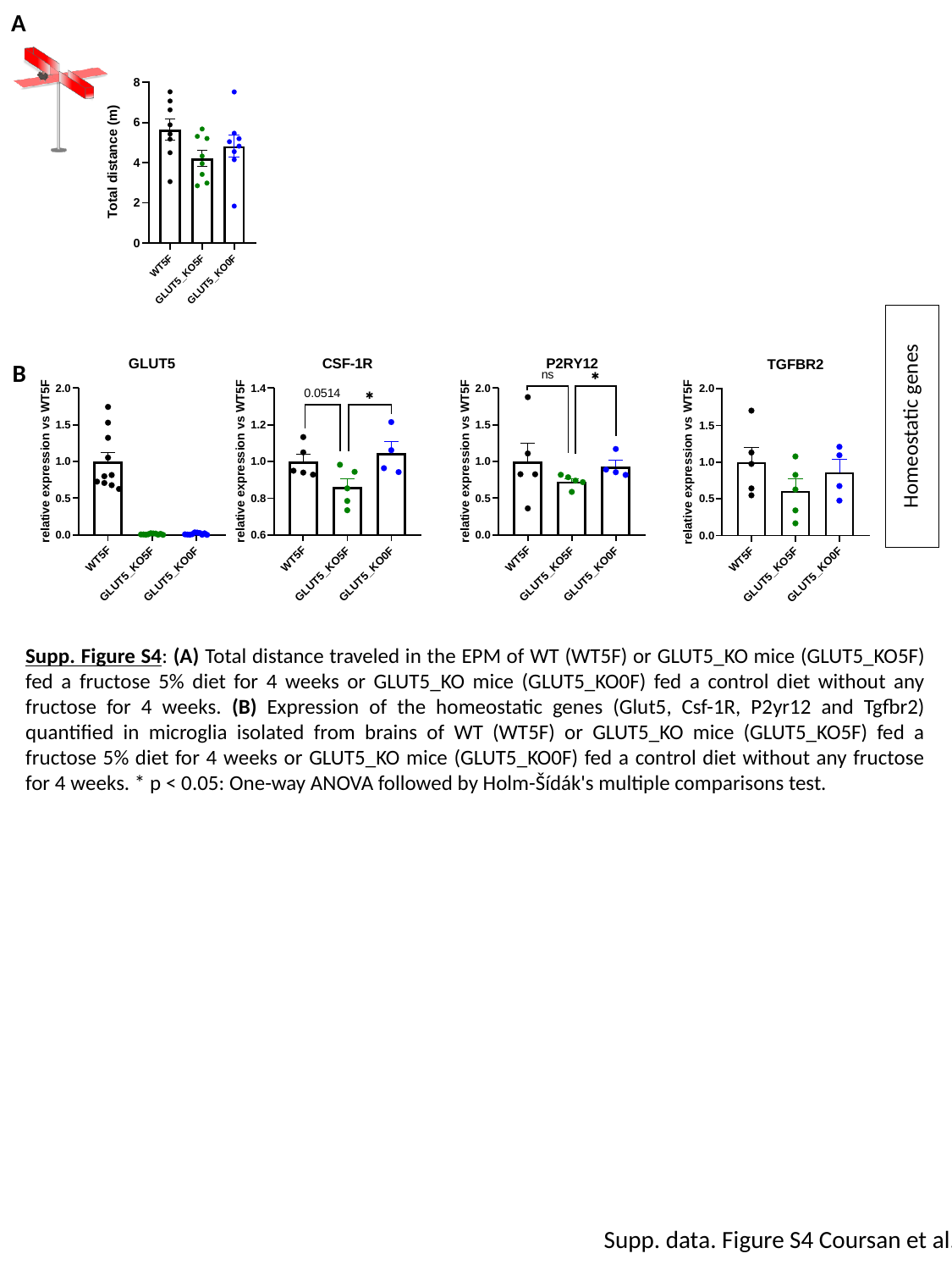

A
B
Homeostatic genes
Supp. Figure S4: (A) Total distance traveled in the EPM of WT (WT5F) or GLUT5_KO mice (GLUT5_KO5F) fed a fructose 5% diet for 4 weeks or GLUT5_KO mice (GLUT5_KO0F) fed a control diet without any fructose for 4 weeks. (B) Expression of the homeostatic genes (Glut5, Csf-1R, P2yr12 and Tgfbr2) quantified in microglia isolated from brains of WT (WT5F) or GLUT5_KO mice (GLUT5_KO5F) fed a fructose 5% diet for 4 weeks or GLUT5_KO mice (GLUT5_KO0F) fed a control diet without any fructose for 4 weeks. * p < 0.05: One-way ANOVA followed by Holm-Šídák's multiple comparisons test.
Supp. data. Figure S4 Coursan et al.

### Slide 5
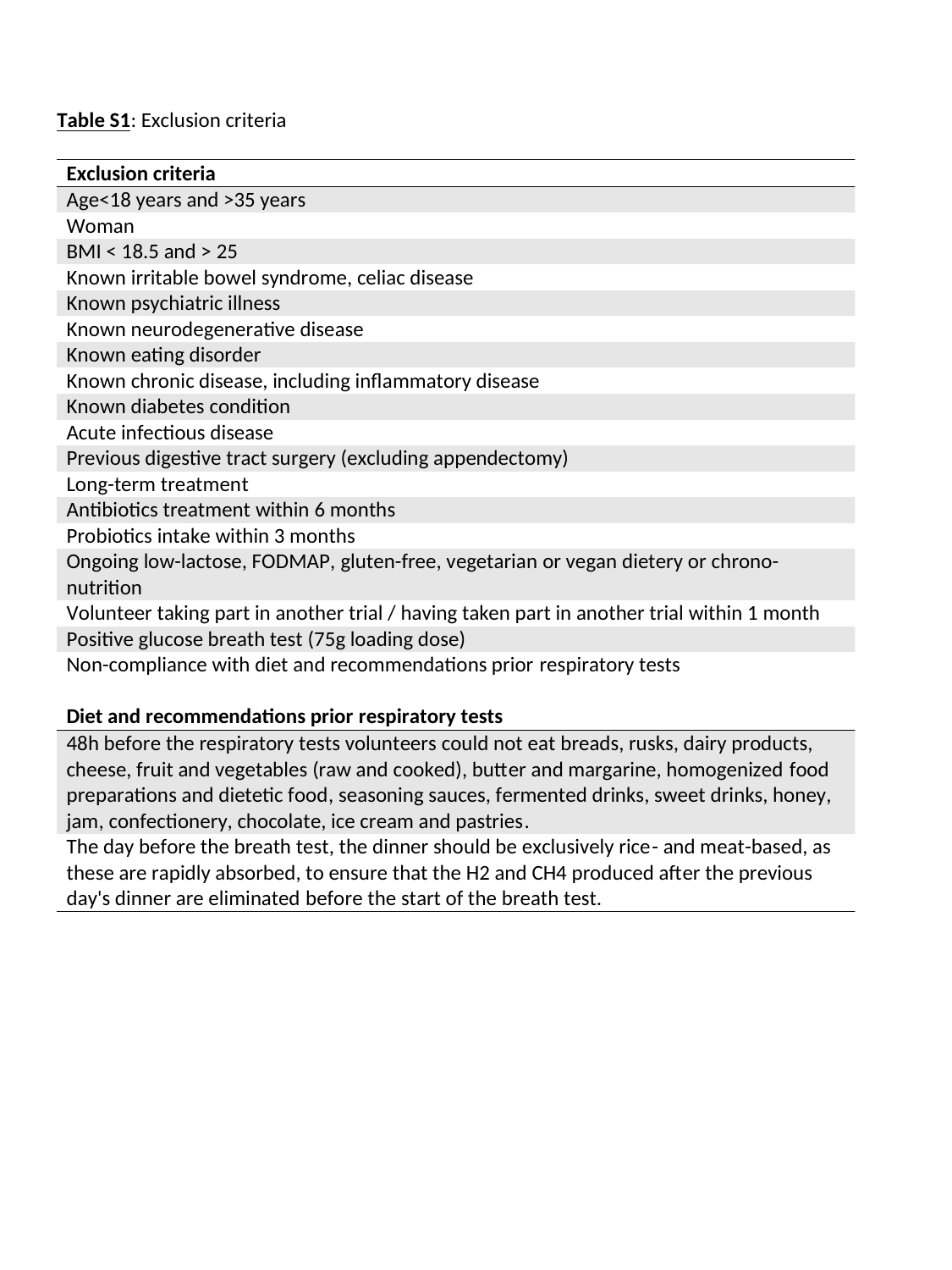

### Slide 6
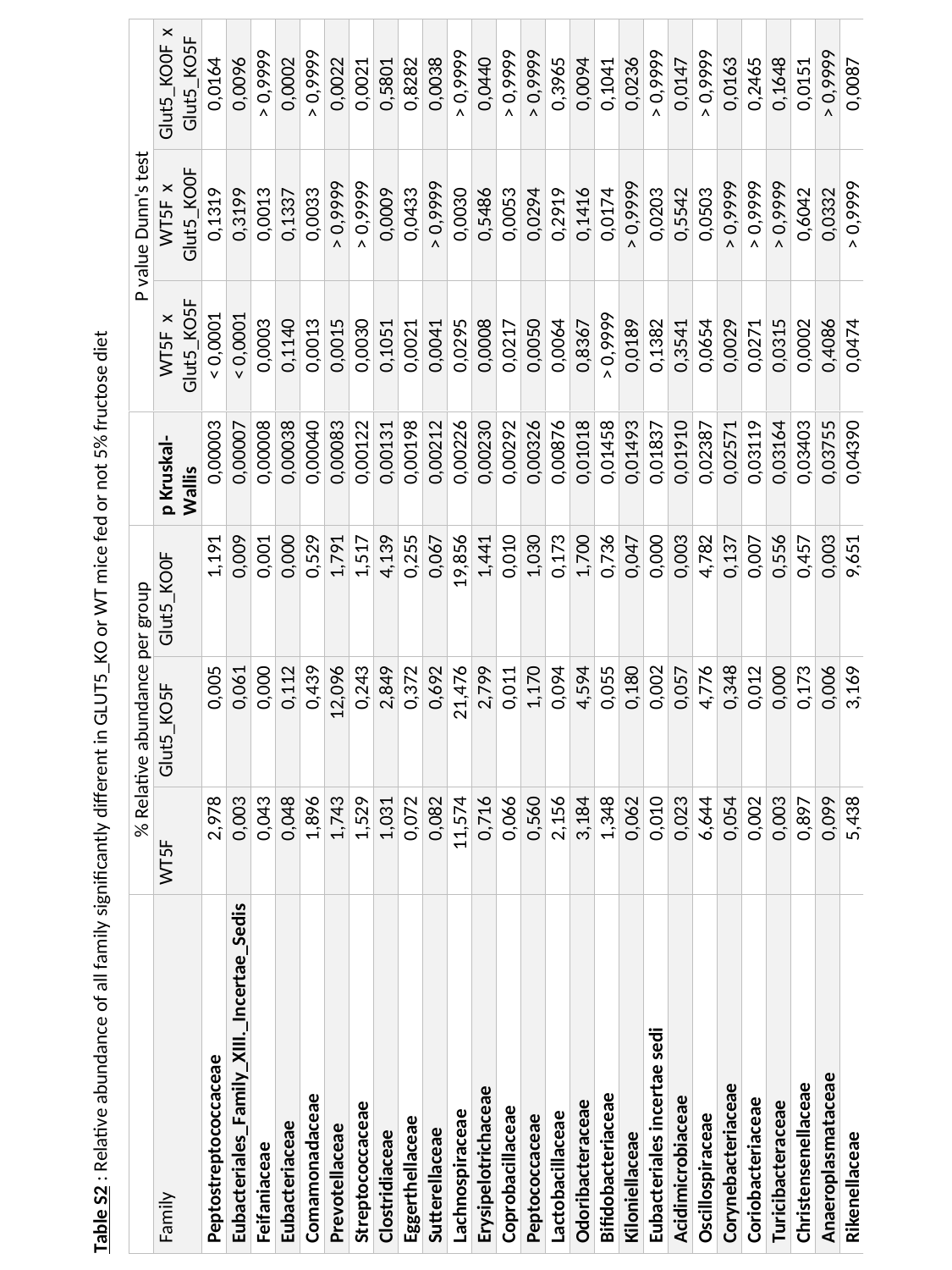
